# Consequences of resource constraint on stochastic gene regulation

**DOI:** 10.1101/2025.04.02.646804

**Authors:** Utkarsh Singh Solanki, Abhyudai Singh, Abhilash Patel

## Abstract

In this contribution, we systematically investigate how intracellular constraints on resources impact stochastic gene expression and regulation. We first consider a model of a single gene with discrete integer-valued mRNA and protein copy numbers that evolved stochastically based on probability occurrences of biochemical reactions. The resource constraints are imposed by considering a finite number of ribosomes binding to mRNAs to form a translation complex, and the complex dissociates to give back a free ribosome and a protein molecule. Analytical analysis reveals that ribosomal constraints reduce the magnitude of stochastic fluctuations in protein copy numbers, and also lead to lower statistical single-cell concordance between mRNA and protein levels of the same gene. We also identify parameter regimes where copy-number fluctuations become sub-Poisson – less variation than expected from a Poisson distribution. Considering fast ribosomal binding/unbinding to mRNAs, we also develop a reduced stochastic model that faithfully captures the statistical fluctuation of the system. Finally, we extend the model to consider an incoherent feedforward loop, and our results show noise minimization by resource constraints in gene regulation.

## 1. INTRODUCTION

Synthetic biology has taken a significant leap forward in designing biomolecular systems with desired phenotypes, facilitated by advances in control engineering [1]–[3]. A fundamental objective of synthetic biology is to achieve precise control of gene expression, an inherently complex biological process by which genetic information encoded within DNA is converted into functional molecules such as mRNA and proteins. Precise control of gene expression can establish numerous promising applications, including therapeutics (e.g., engineered cells for targeted drug delivery [4], personalized medicine [5], and gene therapies [6]), materials science (e.g., biofabrication of advanced biomaterials [7]), renewable energy (e.g., biofuels and bioreactors [8]), and environmental remediation (e.g., biosensors for pollution detection and engineered microbes for waste management [9]). However, reliable implementation of such applications remains challenging due to inherent biological noise arising from stochastic fluctuations in molecular interactions and reaction kinetics. This randomness introduces variability in gene expression levels and leads to substantial differences in behavior even among genetically identical cells within identical environmental contexts, even under tightly regulated conditions [10]–[18]. This variability or expression noise has been shown to have tremendous consequences for the emergence of rare drug tolerant cancer and bacterial cells within otherwise isogenic populations [19]–[28]. Thus, effectively managing these stochastic variations remains a critical hurdle in developing predictable and robust synthetic biological systems.

In engineered biomolecular circuits, the availability of cellular resources, such as ribosomes, RNA polymerases, and metabolic precursors, is essential for proper function. In particular, ribosomes are a critical resource required for protein synthesis, as they bind to mRNA molecules to form translation complexes. However, the intracellular pool of ribosomes is limited, creating competition among different genes being simultaneously expressed within a cell [29]. This competition can cause unintended interactions, leading to resource constraints and significantly altering gene expression dynamics [30]–[32]. Such resource limitations may amplify or attenuate stochastic fluctuations, thereby influencing the overall robustness and predictability of synthetic circuits. Understanding how these finite cellular resources shape the stochastic behavior of gene regulatory networks remains crucial for the rational design of reliable synthetic biological systems.

In this study, we investigated the effects of resource constraints, specifically the limitation of available ribosomes, on stochastic gene expression. We developed mathematical models that explicitly incorporate ribosome availability as a constraint and analyze the resulting gene expression dynamics using stochastic simulations based on chemical master equations. First, we considered a simple gene regulatory system to investigate how constraints impact fluctuations in mRNA and protein levels, using Fano factor. By comparing constrained and unconstrained scenarios, we assessed the effects of limited ribosomal resources on gene expression variability, capturing shifts in stochastic behavior. We found that constraint can reduce the stochasticity, possibly due to unintended proportional-like feedback. Subsequently, we extended our analysis to more complex gene regulatory systems, such as incoherent feedforward loop, to understand how resource limitations influence the stochastic performance and robustness in a larger regulatory motifs. Our results highlight critical parameter regimes where resource constraints lead to noise reduction, thereby contributing valuable insights toward robust synthetic biological circuit design.

## II. Model formulation of resource constraints in gene expression

To study the effects of resource constraints, we consider a set of co-expressing genes ***X*** = {*X*_1_, *X*_2_, …, *X*_*n*_} in a biomolecular system. Each gene *X*_*i*_ transcribes mRNA *X*_*mi*_ constitutively at a rate *α*_*mi*_, while the mRNA is degraded and diluted effectively at a rate *γ*_*mi*_. To translate, the mRNA *X*_*mi*_ binds to ribosomes *R* at a rate *k*_*bi*_, forming a complex *C*_*i*_, which can dissociate at a rate *k*_*ui*_. The translation complex *C*_*i*_ synthesizes the corresponding protein *X*_*pi*_ at a rate *α*_*pi*_, releasing both the mRNA and the ribosome back into the cellular pool for reuse. The protein *X*_*pi*_ subsequently undergoes dilution and degradation at an effective rate *γ*_*pi*_. The chemical reactions describing this process are given by,

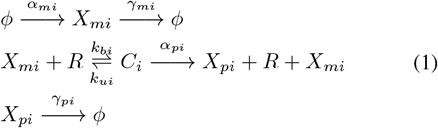

and makes a key assumption that mRNA binding to ribosome protects it from degradation.

In a deterministic framework, the set of chemical reactions can be modeled, using mass action law, as,

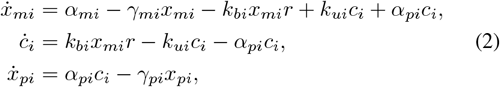

where, *x*_*mi*_, *c*_*i*_, *x*_*pi*_, *r* represent concentrations of *X*_*mi*_, *C*_*i*_, *X*_*pi*_ and resources *R* respectively for genes *i* = 1, … *n*.

Multiple genes can compete for limited cellular resources such as ribosomes, as illustrated in Fig. 1. This competition can lead to resource constraints, as the availability of cellular resources is inherently finite. Furthermore, synthetic biomolecular systems rely on these cellular resources to perform intended functions, and excessive resource consumption can impose a metabolic burden on the host cell. To capture the effects of these constraints, we define the total finite cellular resources *r*_*T*_ as the sum of freely available resources *r* and ribosomes bound to mRNA molecules in the form of translation complexes *c*_*i*_,

**Fig. 1.**
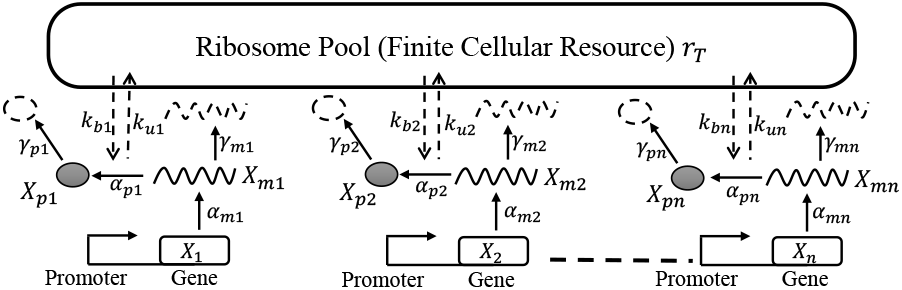
Schematic for co-expression of *n* genes sharing finite cellular resources.

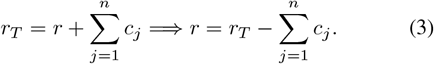

Here, *n* represents the number of genes. In a cell, when resources *r*_*T*_ are abundant, such that the removal of *r* due to 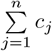 is negligible, gene expression becomes unconstrained, leading to *r* = *r*_*T*_ [31]. Alternatively, the system can be considered unconstrained when the total amount of complexes is sufficiently small, rendering their contribution negligible in comparison to the total resource pool *r*_*T*_. This can be achieved with a parametric combination such as large value of *α*_*pi*_ or *k*_*ui*_. It has been noted that these resource limitations can significantly alter the dynamic behavior of gene expression, demonstrated in [31], by effectively introducing additional feedback layers into the system. This is also evident from Eq. (2), where *c*_*i*_ feeds back into the mRNA dynamics.

## III Resource constraints in expression of a single gene

We first try to quantify the impact of resource constraints for a single gene, where multiple mRNAs of the same gene compete for ribosomes. In this case, the system of chemical reactions (1) takes the form (for simplicity, the index *i* is dropped for a single gene),

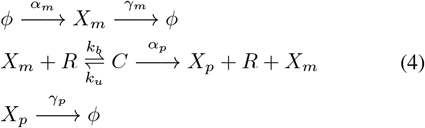

### A. Deterministic formulation for ribosome competition at a single gene

Mass action kinetics for (4) results in the following nonlinear system of ordinary differential equations

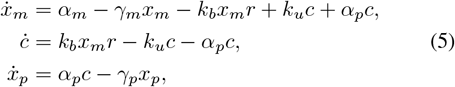

that has unique asymptotically stable equilibrium point [33],

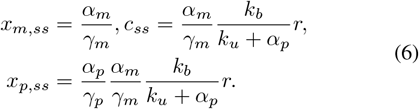

Note that the steady-state level of the free (not bound to ribosomes) mRNA *x*_*m,ss*_ only depends on the transcription and degradation rate *α*_*m*_ and *γ*_*m*_, respectively, and independent of ribosome binding/unbinding/translation kinetics. This result is reminiscent of results on decoy binding of proteins, where binding protects the protein from active degradation [34]–[36]. Using the fact that *r* + *c*_*ss*_ = *r*_*T*_ we have the free ribosome level as

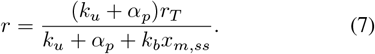

From the expression for *r*, we observe that as the steadystate mRNA concentration *x*_*m,ss*_ increases, the pool of available ribosomes *r* for translation decreases. The binding dynamics between ribosomes and mRNA influence ribosome availability: increasing the binding rate *k*_*b*_ reduces the free ribosome pool, while increasing the unbinding rate *k*_*u*_ replenishes the available ribosomes, although this effect saturates at high values of *k*_*u*_. As we increase the translation rate *α*_*p*_, the available ribosome *r* tends toward *r*_*T*_, and at *α*_*p*_ → ∞, *r* = *r*_*T*_ leads to a purely unconstrained system. Since resource availability is a critical cellular parameter, it can modulate the magnitude of intrinsic stochastic fluctuations arising from biochemical reactions operating at low copy numbers. We next investigate the stochastic formulation of the system of bioreactions (4), with discrete integer copy number of species that evolve with random occurrences of individual reactions [37]–[39].

### B. Analysis of expression heterogeneity via simulations

To illustrate these concepts, we begin with a simplified model of single-gene expression (Fig. 2(a), inset), which serves as the foundation for understanding how resource constraints influence stochastic gene regulation. In this system, transcription, translation, and degradation processes are modeled as discrete stochastic events, with each reaction governed by a propensity function that determines the probability of its occurrence per unit time. The set of reaction propensities and corresponding state transitions for this single-gene system are summarized in Table I, and this describes the time evolution of the integer-valued population counts ***x***_***m***_, ***c, x***_***p***_ ∈ {0, 1, …} for species *X*_*m*_, *C* and *X*_*p*_ in (4). The level of the free ribosomes *r* is *r*_*T*_ − ***c***. For simplicity we assume the total ribosome count *r*_*T*_ to be fixed, but in reality this itself can be a stochastic process contributing to “extrinsic noise” in gene expression [40], [41]. Please note the bold notation used for species counts here as they are random processes.

**TABLE 1.** Propensities for population reset for corresponding reactions for single gene expression.

| Reaction | Population Reset | Propensity |
| --- | --- | --- |
| $\phi \rightarrow X_m$ | $x_m \rightarrow x_m + 1$ | $\alpha_m$ |
| $X_m \rightarrow \phi$ | $x_m \rightarrow x_m - 1$ | $\gamma_m x_m$ |
| $X_m + R \rightarrow C$ | $c \rightarrow c + 1,$<br>$x_m \rightarrow x_m - 1$ | $k_b (r_T - c) x_m$ |
| $C \rightarrow X_m + R$ | $c \rightarrow c - 1,$<br>$x_m \rightarrow x_m + 1$ | $k_u c$ |
| $C \rightarrow X_p + X_m + R$ | $x_p \rightarrow x_p + 1,$<br>$c \rightarrow c - 1,$<br>$x_m \rightarrow x_m + 1$ | $\alpha_p c$ |
| $X_p \rightarrow \phi$ | $x_p \rightarrow x_p - 1$ | $\gamma_p x_p$ |

**Fig. 2.**
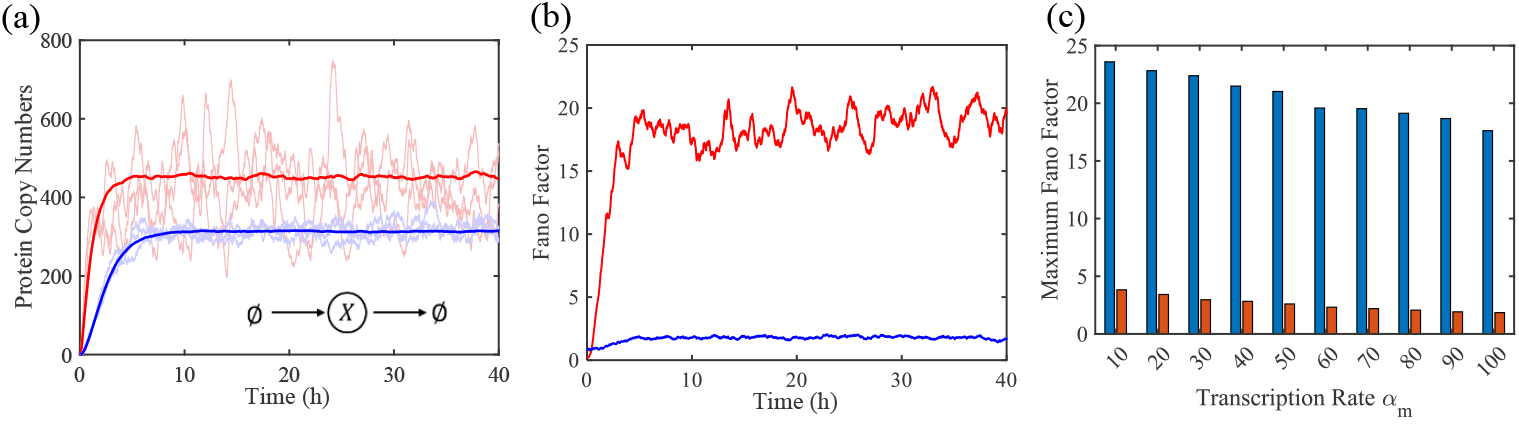
Simulations for simple gene expression (inset). The key parameters for the model used were considered as follows: total ribosome numbers assumed to be *r*_*T*_ = 50 molecules per cell and rates *k*_*b*_ = 5 *h*^*−*1^, *k*_*u*_ = 50 *h*^*−*1^, *γ*_*m*_ = 10 *h*^*−*1^, *α*_*m*_ = 20 *h*^*−*1^, and *γ*_*p*_ = 1 *h*^*−*1^. All these parameters are scaled with respect to the protein decay rate. The simulation was performed for 500 trajectories. (a) Mean of ***x***_*p*_ counts in a resource constraint scenario due to *α*_*p*_ = 10 (blue solid line) and in a scenario without resource constraint (red solid line) due to *α*_*p*_ = 400 for 500 trajectories. A few sample trajectories from constraint and without constraints simulations are presented transparently in the respective colors. (b) Fano factor, variance over mean for 500 trajectories, of ***x***_*p*_ with resource constraint (blue line, *α*_*p*_ = 10) and without resource constraint (red line, *α*_*p*_ = 400) (c) The maximum value of Fano factor of ***x***_*p*_ across time from the simulations, indicated as Maximum Fano Factor, for change in the mRNA transcription rate *α*_*m*_ from 10 to 100. The blue bar represents the without constraint *α*_*p*_ = 400 and orange bar represents with constraint *α*_*p*_ = 10.

These propensities form the basis for simulating the time evolution of the system using the Stochastic Simulation Algorithm (SSA), also known as the Gillespie algorithm [42]. Unlike deterministic models that rely on average behavior, SSA captures the intrinsic noise by simulating each individual reaction event, allowing us to study the full distribution of gene expression levels across many stochastic trajectories. This approach enables us to directly quantify variability in protein expression under both resource-constrained and unconstrained scenarios. The stochasticity in protein expression can be quantitatively assessed using the Fano factor, defined as the variance-to-mean ratio of protein copy numbers. This metric provides a normalized measure of noise and allows direct comparison across different conditions or circuit configurations. In the case of a purely Poissonian process—where each event occurs independently and with constant probability—the variance equals the mean, resulting in a Fano factor of one [12], [40]. A Fano factor greater than one typically signifies super-Poissonian noise, which may arise from transcriptional bursts, resource fluctuations, or feedback delays [43]. In contrast, a Fano factor less than one (sub-Poissonian noise) suggests noise suppression, often due to buffering mechanisms, effective negative feed-back, or existence of multiple rate limiting steps in the transcription and mRNA degradation pathways [44]. In our context, introducing ribosomal resource constraints creates an emergent feedback-like behavior: as mRNA abundance increases, the competition for ribosomes intensifies, naturally limiting translation rates. This self-limiting mechanism can suppress fluctuations and result in sub-Poissonian statistics.

To compare protein levels and the corresponding Fano factor for single-gene expression under resource-constrained and unconstrained conditions, stochastic simulations were performed using the Gillespie algorithm, as shown in Fig. 2. The simulations are based on the model described in Eq. (5). For a single gene, the propensity for mRNA transcription is *α*_*m*_, while mRNA degradation follows a propensity of *γ*_*m*_***x***_***m***_. Ribosome binding to mRNA occurs with a propensity of *k*_*b*_(*r*_*T*_ − ***c***)***x***_***m***_, and unbinding occurs with a propensity of *k*_*u*_***c***, where *r*_*T*_ represents the pool of ribosomes and ***c*** is the ribosome-mRNA complex. Protein synthesis proceeds at a rate *α*_*p*_***c***, and protein degradation follows the propensity *γ*_*p*_***x***_***p***_. The complete set of reaction propensities and their associated population updates are provided in Table I. These simulations allow us to assess how resource availability influences both the mean expression levels.

For a high translation rate *α*_*p*_, the protein copy number reaches a higher steady state, with significantly large variability across trajectories (Fig. 2(a)). In contrast, when available resources are decreased by decreasing the translation rate *α*_*p*_, the mean protein level along with variability decreases, possibly due to unintended competitive interactions for ribosomes. This stochastic aspect is quantitatively captured by the Fano factor (variance-to-mean ratio), as shown in Fig. 2(b). Under conditions of abundant resource availability, the Fano factor exceeds one, indicating super-Poissonian noise—likely a consequence of unregulated resources enabling burstier translation dynamics. In contrast, under limited resource conditions, the Fano factor decreases significantly, reflecting a suppression of gene expression noise. To further investigate, we examined the factors that contribute to increased resource demand and consequently impose stronger constraints on the system. We increased the resource consumption by increasing the transcription rate (Fig. 2(c)). Increasing the transcription rate, increases the mRNA copy numbers, which ultimately decreases the available ribosomes, as shown in Eq. (7). We considered the maximum Fano factor as a metric because this will capture the worst-case scenario of unpredictability. We noted that lower transcription rates result in higher Fano factors, and as the transcription rate increases, the maximum Fano factor decreases, irrespective of the translation rate. This indicates the noise attenuation is due to the resource constraint.

### C. Analytical quantification of protein count statistics with resource constraints

To develop closed-form formulas for the magnitude of statistical fluctuations in gene product copy numbers we use the framework of moment dynamics for chemical reactions [45]. For example, consider the statistical moment 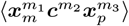, where ⟨.⟩ is the expected value operator and *m*_1_, *m*_2_, *m*_3_ are appropriately chosen nonnegative integers. Then, the time evolution of this moment is given by [45]

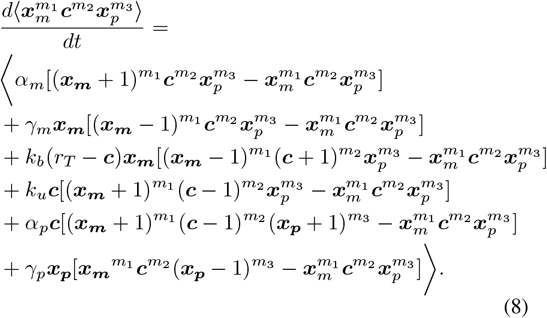

Using this, the time evolution of the means is given by

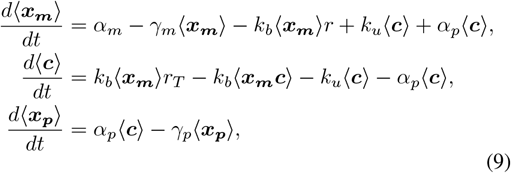

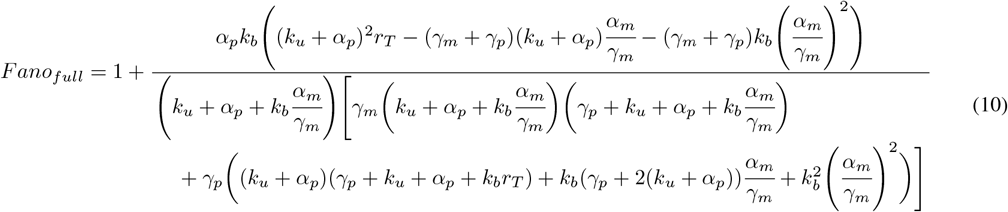

a system analogous to mass action kinetics (4). Notice the presence of the second-order moment ⟨***x***_*m*_***c⟩*** which makes the system of differential equations not solvable, and generally moment closure schemes are employed to obtain approximate solutions [46]–[52].

To obtain an approximate analytical solution of the steady-state moments, we here use the Linear Noise Approximation (LNA) [53], [54] that works by first linearizing the nonlinear propensities around the solution of the mass action kinetics and then deriving the moment dynamics with the *linearized* propensities. Note from Table I, the nonlinearity in the propensity for the third reaction that is approximated by

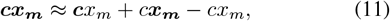

where the unbold *x*_*m*_, *c* are solutions to Eq. (5). In this LNA approximation, the time evolution of the statistical means reduces to mass action kinetics, and the time evolution of the second-order moments results in a “closed” system of differential equations [54], [55] that we then solve to obtain the steady-state moments. The steady-state Fano factor of the protein copy number obtained from LNA is shown in Eq. (10) at the top of the next page.

In Fig. 3, we plot the steady-state Fano factor Eq. (10) for decreasing mRNA translation rate. This analysis shows a decrease in Fano factor with increasing ribosome competition among mRNA captured by reducing *α*_*p*_. The Fano factor approaches a value of one as *α*_*p*_ → 0, and approaches a value of

**Fig. 3.**
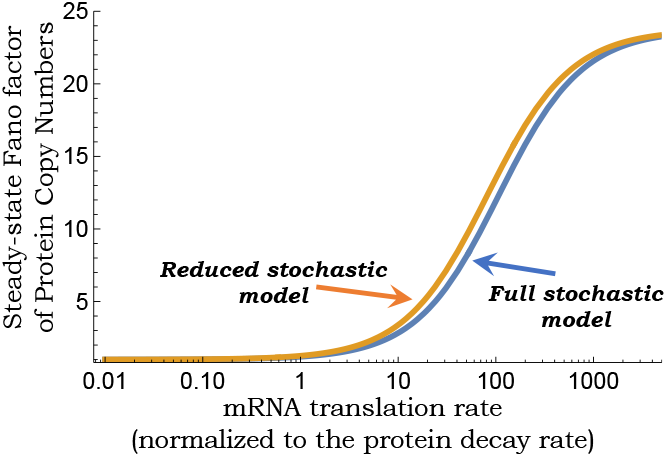
The steady-state Fano factor for quantifying fluctuations in the protein counts in the full model in Table I (blue line) and the reduced model in Table II (yellow line) as obtained by using the Linear Noise Approximation. The Fano factor is plotted as a function of the mRNA translation rate *α*_*p*_ with total ribosome numbers assumed to be *r*_*T*_ = 50 molecules and rates *k*_*b*_ = 5 *h*^*−*1^, *k*_*u*_ = 50 *h*^*−*1^, *γ*_*m*_ = 10 *h*^*−*1^, *α*_*m*_ = 20 *h*^*−*1^, and *γ*_*p*_ = 1 *h*^*−*1^.

**TABLE 2.**
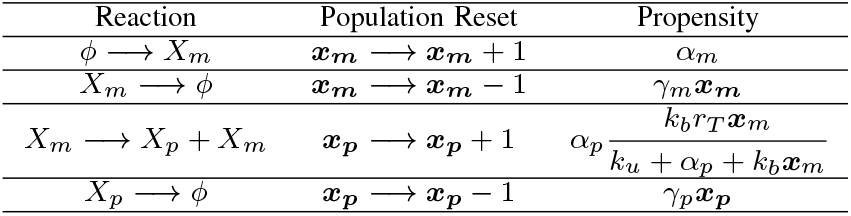
Propensities for population reset for corresponding reactions for reduced single gene expression.

| Reaction | Population Reset | Propensity |
| --- | --- | --- |
| $\phi \rightarrow X_m$ | $\mathbf{x}_m \rightarrow \mathbf{x}_m + 1$ | $\alpha_m$ |
| $X_m \rightarrow \phi$ | $\mathbf{x}_m \rightarrow \mathbf{x}_m - 1$ | $\gamma_m \mathbf{x}_m$ |
| $X_m \rightarrow X_p + X_m$ | $\mathbf{x}_p \rightarrow \mathbf{x}_p + 1$ | $\alpha_p \frac{k_b r_T \mathbf{x}_m}{k_u + \alpha_p + k_b \mathbf{x}_m}$ |
| $X_p \rightarrow \phi$ | $\mathbf{x}_p \rightarrow \mathbf{x}_p - 1$ | $\gamma_p \mathbf{x}_p$ |

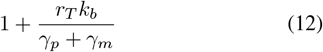

for *α*_*p*_ *→ ∞*, as obtained in classical model of stochastic gene expression with no constraints on ribosome availability [56], [57]. Interestingly, our results show that when *k*_*u*_ = 0, then the Fano factor can reduce below one (sub-Poissonian fluctuations) for intermediate values of *α*_*p*_ (Fig. 4).

**Fig. 4.**
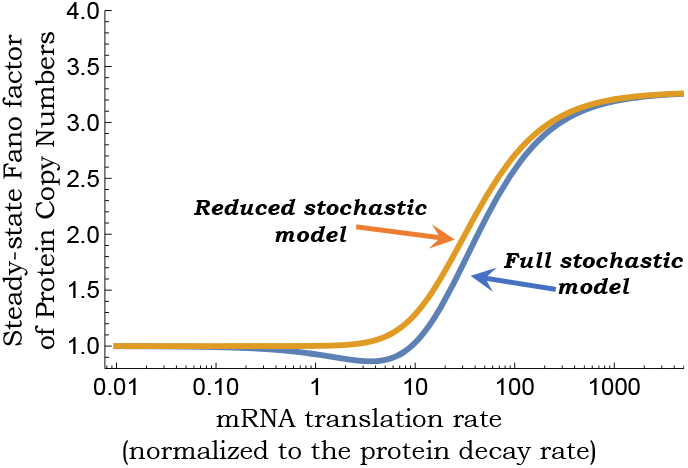
The steady-state Fano factor for quantifying fluctuations in the protein counts for full (Table I) and reduced (Table II) models as obtained by using the Linear Noise Approximation. All rates are in Fig. 3 except *k*_*u*_ = 0.

### D. Reduced stochastic model capturing resource constraints

In the limit of fast ribosome binding and unbinding, we consider a reduced two-dimensional model where the stochastic dynamics of the complex is eliminated, and protein synthesis occurs with the nonlinear propensity

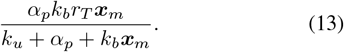

This creates the feedback alluded to earlier, where high levels of ***x***_*m*_ are compensated for by reduced protein synthesis. The reduced model is illustrated in Table II, and applying the LNA to the reduced model results in the steady-state protein Fano factor

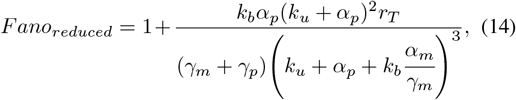

where one can clearly see the limits

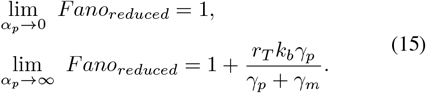

The Fano factor from the reduced model faithfully captures the behavior of the full model (Fig. 3), but fails to capture the sub-Poisson regime that arises for slow kinetics of ribosomes unbinding and translation. The reduced model also predicts the normalized steady-state covariance between mRNA and its protein

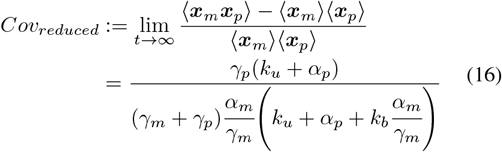

where in the limits of rapid translation

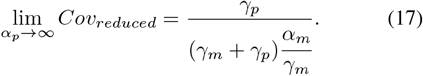

With increasing competition for ribosomes captured by decreasing *α*_*p*_, *Cov*_*reduced*_ reduces predicting lesser concordance between the copy number of mRNA and its protein at the single-cell level.

### IV. Resource constraints in complex gene regulatory networks

To investigate the impact of resource constraints on larger gene regulatory networks, we consider the incoherent feed-forward loop [58], which plays an essential role in achieving homeostasis. In an incoherent feedforward loop (IFFL), an input signal regulates a target gene through two distinct pathways, direct activation and indirect repression, which oppose each other (Fig. 5(a), inset). In this incoherent feedforward loop, the input signal *u* directly activates the expression of both the genes *X* and *Y*, and simultaneously, the gene *X* encodes a repressor protein *X*_*p*_, which represses the gene *Y*. This regulatory configuration produces pulse-like dynamics [59].

**Fig. 5.**
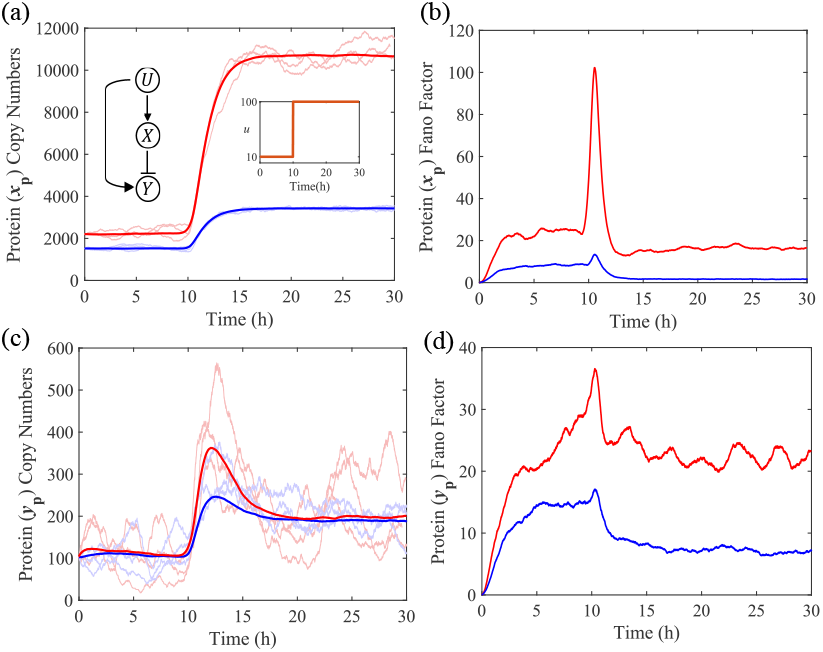
Simulations for incoherent feedforward loop for a step input of *u* from 10 to 100 at time = 10 *h* (inset, a). The key parameters for the model used were considered as follows: the rates are *α*_*mx*_ = *α*_*my*_ = 500 *h*^*−*1^, *γ*_*mx*_ = *γ*_*my*_ = 1 *h*^*−*1^, *k*_1_ = *k*_2_ = 100, *k*_*bx*_ = *k*_*by*_ = 1 *h*^*−*1^, *k*_*ux*_ = *k*_*uy*_ = 1 *h*^*−*1^, *γ*_*px*_ = *γ*_*py*_ = 1 *h*^*−*1^, and the total ribosome per cell *r*_*T*_ is given as 50 molecules. All these parameters are scaled with respect to the protein decay rate. (a) Mean of ***x***_*p*_ copy numbers, where blue solid line represents low translation rate *α*_*px*_ = *α*_*py*_ = 100 *h*^*−*1^ and red solid line represents high translation rate *α*_*px*_ = *α*_*py*_ = 1500 *h*^*−*1^, for 500 trajectories for protein *X*_*p*_. Sample trajectories for low resources and high resources are presented transparently in the respective colors, (b) Fano factor, variance over mean for 500 trajectories, of ***x***_*p*_ with low translation rate is represented as blue line and with high translation rate is represented as red line, (c) Mean of ***y***_*p*_ copy numbers, where blue solid line represents low translation rate *α*_*px*_ = *α*_*py*_ = 100 *h*^*−*1^ and red solid line represents high translation rate *α*_*px*_ = *α*_*py*_ = 1500 *h*^*−*1^, for 500 trajectories for protein *Y*_*p*_. Sample trajectories for low resources and high resources are presented transparently in the respective colors, (d) Fano factor, variance over mean for 500 trajectories, of ***y***_*p*_ with low translation rate is represented as blue line and with high translation rate is represented as red line.

To model the incoherent feedforward loop using the stochastic simulation algorithm (SSA), we explicitly consider all relevant biochemical reactions. The probability per unit time for the reactions as a stochastic system can be written using propensity functions as stated in Table III. The mRNA *X*_*m*_ is transcribed at the rate *α*_*mx*_*f* (*u*) where 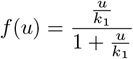 and *u* is a step signal and *k*_1_ is the dissociation constant, and this mRNA is degraded and diluted effectively at the rate *γ*_*mx*_. The mRNA *X*_*m*_ binds to ribosomes *R* at the rate *k*_*bx*_ to form the translation complex *C*_*x*_, which dissociates at the rate *k*_*ux*_. The translation complex *C*_*x*_ synthesizes protein *X*_*p*_ at the rate *α*_*px*_, releasing free mRNA and ribosomes. The protein *X*_*p*_ undergoes dilution or degradation at the rate *γ*_*px*_. Similarly, mRNA *Y*_*m*_ is transcribed at a rate *α*_*my*_*g*(*u*, ***x***_***p***_), where 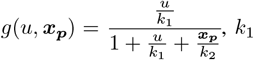 is dissociation constant of *u*, and *k*_2_ is the dissociation constant of *X*_*p*_. The mRNA *Y*_*m*_ is degraded and diluted combinedly at the rate *γ*_*my*_. The mRNA *Y*_*m*_ binds to ribosomes *R* at the rate *k*_*by*_ to form the translation complex *C*_*y*_, dissociating at the rate *k*_*uy*_. We assume reaction rates for gene *X* and *Y* expressions to be the same. This complex *C*_*y*_ produces protein *Y*_*p*_ at the rate *α*_*py*_, again releasing free mRNA and ribosomes for reuse. Protein *Y*_*p*_ also undergoes dilution or degradation at the rate *γ*_*py*_.

**TABLE 3.** Propensities for population reset for corresponding reactions for Incoherent Feedforward loop.

| Reaction | Population Reset | Propensity |
| --- | --- | --- |
| $\phi \xrightarrow{f(u)} X_m$ | $x_m \rightarrow x_m + 1$ | $\alpha_{mx} f(u)$ |
| $X_m \rightarrow \phi$ | $x_m \rightarrow x_m - 1$ | $\gamma_{mx} x_m$ |
| $X_m + R \rightarrow C_X$ | $c_x \rightarrow c_x + 1,$<br>$x_m \rightarrow x_m - 1$ | $k_{bx} x_m (r_T - c_x - c_y)$ |
| $C_X \rightarrow X_m + R$ | $c_x \rightarrow c_x - 1,$<br>$x_m \rightarrow x_m + 1$ | $k_{ux} c_x$ |
| $C_X \rightarrow X_p + X_m + R$ | $x_p \rightarrow x_p + 1,$<br>$c_x \rightarrow c_x - 1,$<br>$x_m \rightarrow x_m + 1$ | $\alpha_{px} c_x$ |
| $X_p \rightarrow \phi$ | $x_p \rightarrow x_p - 1$ | $\gamma_{px} x_p$ |
| $\phi \xrightarrow{g(u, x_p)} Y_m$ | $y_m \rightarrow y_m + 1$ | $\alpha_{my} g(u, x_p)$ |
| $Y_m \rightarrow \phi$ | $y_m \rightarrow y_m - 1$ | $\gamma_{my} y_m$ |
| $Y_m + R \rightarrow C_Y$ | $c_y \rightarrow c_y + 1,$<br>$y_m \rightarrow y_m - 1$ | $k_{by} y_m (r_T - c_x - c_y)$ |
| $C_Y \rightarrow Y_m + R$ | $c_y \rightarrow c_y - 1,$<br>$y_m \rightarrow y_m + 1$ | $k_{uy} c_y$ |
| $C_Y \rightarrow Y_p + Y_m + R$ | $y_p \rightarrow y_p + 1,$<br>$c_y \rightarrow c_y - 1,$<br>$y_m \rightarrow y_m + 1$ | $\alpha_{py} c_y$ |
| $Y_p \rightarrow \phi$ | $y_p \rightarrow y_p - 1$ | $\gamma_{py} y_p$ |

$f(u) = \frac{u/k_1}{1+u/k_1}$ is an activating function, $g(u, x_p) = \frac{u/k_1}{1+x_p/k_2+u/k_1}$ is a combinatorial regulation function. They scale the transcription rate.

In a deterministic framework, the chemical reactions can be modeled using mass action law as,

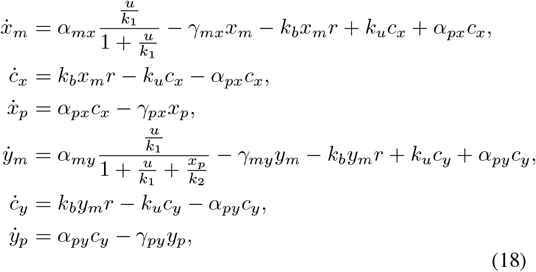

where, *x*_*m*_, *c*_*x*_, *x*_*p*_, *y*_*m*_, *c*_*y*_, *y*_*p*_, *r* represent concentrations of *X*_*m*_, *C*_*x*_, *X*_*p*_, *Y*_*m*_, *C*_*y*_, *Y*_*p*_ and *R* respectively.

Similar to the single-gene system, resource availability can be modulated by varying the translation rate. At a high value of the translation rate, the number of free ribosomes in the cell is sufficient to approximate an unconstrained system. Conversely, at lower translation rates, resource availability drops significantly, introducing a constraint on gene expression within the IFFL.

The dynamics of protein concentration and the corresponding Fano factor for the incoherent feedforward loop are illustrated using stochastic simulations (Gillespie algorithm) in Fig. 5, based on the model described in Eq. (18). The protein copy number ***x***_*p*_ is substantially reduced when the available resource is limited due to the low translation rate, compared to the scenario of a higher translation rate (Fig. 5(a)). This is expected as the limited resource creates a negative interaction among competitive species, as seen in single gene expression also. The Fano factor (variance-to-mean ratio) for protein *X*_*p*_ is notably lower under resource constraint conditions. Without constraints, the Fano factor stabilizes at a significantly higher value, indicating that the unconstrained scenario is more susceptible to stochastic variability (Fig. 5(b)). The presence of resource constraints reduces the amplitude of the protein pulse, indicating a direct impact of ribosomal limitation on the performance of the system (Fig. 5(c)). It is interesting to note that the resource constraint does not change the steady-state level or alter the adaptation capability of IFFL. Similar to the protein *X*_*p*_, with resource constraints, the fluctuations in protein *Y*_*p*_ are smaller, as evidenced by the lower Fano factor (Fig. 5(d)), reflecting reduced stochastic noise in the system. In both protein concentrations, lower variability and rapid stabilization suggest that resource constraints enhance robustness to noise.

## V. CONCLUSION

In this study, we systematically investigated how intra-cellular resource constraints—specifically ribosome limitations—impact the stochastic dynamics of gene expression. Using a stochastic analysis framework based on moment dynamics and stochastic simulations, we demonstrated that ribosomal competition can significantly suppress variability in protein levels. Starting from a single-gene expression system, we showed that introducing resource constraints can reduce the mean protein output but decrease the Fano factor, indicating a fundamental tradeoff. We further analyzed how gene transcription rates influence noise under constrained and unconstrained conditions. The results revealed that resource constraints act as an intrinsic noise-buffering mechanism, particularly in high-expression regimes. This emergent feedback-like behavior stems from ribosomal saturation, which limits excessive translation and flattens noise amplification across gene copies. Extending the analysis to more complex regulatory motifs, such as the incoherent feed-forward loop, we found that resource constraints consistently contribute to noise minimization. These findings highlight the importance of accounting for intracellular resource limitations in the design of synthetic biological circuits. Overall, our results provide critical insights into how resource-aware circuit design can improve robustness, predictability, and scalability in synthetic biology applications.

